# The dynamics of cell-free DNA from urine and blood after a full marathon

**DOI:** 10.1101/2021.03.06.434188

**Authors:** Yasuhiro Shishikura, Katsuyuki Tokinoya, Yuichi Aita, Nanami Sekine, Takehito Sugasawa, Yasuko Yoshida, Keisei Kosaki, Shota Kumamoto, Keisuke Ishikura, Tomoaki Kuji, Yasushi Kawakami, Yoshiharu Nabekura, Seiji Maeda, Kazuhiro Takekoshi

## Abstract

**Purpose:** Cell-free DNA (cfDNA) has been investigated as a minimally invasive biomarker for many diseases, particularly cancer. An increase in cfDNA has been observed during exercise. Neutrophil extracellular traps (NETs) may be the origin of cfDNA in response to acute exercise, but the mechanisms of generation of cfDNA during exercise remain unclear. In this study we investigated the dynamics of serum and urinary cfDNA levels and determined the relevance of other biomarkers to serum and urinary cfDNA levels and fragment size after a full marathon.

**Methods:** Samples were collected from 23 healthy male subjects. Blood and urine samples were collected before and immediately, two hours, and one day after the full marathon. The measurements included serum and urinary cfDNA, creatine kinase, myoglobin, creatinine, white blood cells, platelets, and lactoferrin from blood, and amylase, albumin, and creatinine from urine.

**Results:** Serum and urinary cfDNA levels increased after a full marathon. Creatine kinase, myoglobin, and creatinine in blood, and albumin and creatinine in urine also increased significantly after a full marathon. Serum cfDNA showed peak values about 180 bp after the full marathon. Values over 1000 bp were present at two hours post-marathon. Urinary cfDNA showed peak values from 35 bp to 50 bp after the full marathon. Values over 1000 bp appeared at Immediately and two hours post marathon.

**Conclusion:** This study revealed that both serum and urinary cfDNA levels transiently increased after a full marathon. In addition, these cfDNA fragment varied in size.

## Introduction

Running has a significant positive effect on the body, improving cardiorespiratory function and preventing and ameliorating various diseases^1^. Recently, the number of people participating in full marathons (42.195 km) has increased in accordance with an increase in the popularity of running^2^. However, previous studies have suggested that completing a full marathon results in systemic damage to various tissues and organs, such as muscle, heart, liver, and kidneys^3–8^. Therefore, there is a need for biomarkers that can identify the conditions within the body during and after endurance exercises.

Cell-free DNA (cfDNA) comprises extracellular free DNA fragments that are released upon stress induction, and that circulate in the body fluids^9^. cfDNA has been examined as a non-invasive diagnostic biomarker for various diseases, including cancer^9,10^. Previous studies have reported that blood cfDNA levels are increased following various exercises^11–16^. Blood cfDNA levels peak immediately after acute exercise and recover rapidly to the baseline level. The typical markers of skeletal muscle damage, such as creatine kinase (CK), are increased several days after exercise^15,16^. Additionally, strenuous exercise causes internal stress that damages leukocytes, injures muscle tissues, and leads to acute inflammatory responses via oxidative stress^3,4,6–8^. Therefore, since cfDNA can be produced in response to several vital reactions, it may be a novel biomarker for responses to exercise including muscle damage, inflammation, and oxidative stress.

cfDNA circulating in the body has several fragment sizes^17^. Small cfDNA fragments of approximately 180 bp in size accumulate in the blood of subjects with a variety of conditions, including pregnancy, cancer, liver or bone marrow transplantation, and systemic lupus erythematosus^18^. Human blood sometimes contains cfDNA fragments larger than 10,000 bp in size, which originate from cell necrosis^19^. Recently, the characteristics of the cfDNA fragment profile have been evaluated in several different cell lines, and it has been found that there are different forms of cfDNA release patterns in every cell line tested^20^. Therefore, measuring the size of cfDNA fragments induced by strenuous exercise can provide insights into the origin and physiological function of cfDNA.

It has been suggested that blood cfDNA induced by exercise is derived not only from apoptosis or necrosis, but also from neutrophil extracellular traps (NETs)^11^. It is possible that increased blood cfDNA levels due to exercise are derived from apoptosis or necrosis-induced skeletal muscle damage in a full marathon, because of the presence of increased muscle damage markers or necrosis after a full marathon^3–5^. Acute exercise causes apoptosis in the skeletal muscle of rats^21^. Atamaniuk et al. showed that blood cfDNA may be derived from leukocytes during an ultra-marathon^12^. Beiter et al. described exercise-induced release of NETs^22^. Recent studies have partly revealed the role of lactoferrin, a component of NETs^23,24^. Lactoferrins inhibit the formation of NETs, possibly by preventing their spread^25^. However, the mechanism of generation of lactoferrin in exercise-induced NETs remains unclear. If a relationship between existing exercise markers and the size of cfDNA fragments induced by exercise can be confirmed, analysis of this relationship may aid in understanding the internal conditions of the body. However, previous studies regarding the effect of cfDNA on exercise have only examined cfDNA in blood, rather than using the non-invasive approach of measuring urinary cfDNA. Resistance exercise increases the production of urinary titin N-terminal fragments and muscle damage markers^26^. Therefore, it is likely that urinary cfDNA changes in concentration and fragment size after a full marathon.

The purpose of this study was to investigate the dynamics of serum and urinary cfDNA levels and fragment sizes after a full marathon. Muscle damage markers, stress hormones, and inflammatory responses known to be affected by exercise were measured to assess the body conditions, and their association with serum and urinary cfDNA levels and fragment sizes after a full marathon was investigated.

## Methods

### Ethical approval

This study was approved by the Ethical Committee of the Faculty of Medicine at the University of Tsukuba (Approval number: 274). All subjects received an explanation and documents in advance regarding the purpose of the experiment, its contents, and safety issues, and indicated their informed consent.

### Subjects

Twenty-six healthy male subjects who undertook aerobic exercise at least twice per week were recruited. The sample size was determined by previous studies^4,27^. The subjects completed the 38th Tsukuba Marathon. Subject characteristics are provided in Table 1. The subjects were instructed not to drink alcohol, to get sufficient sleep, and to avoid binge eating before the experiment. The subjects freely performed warmups and drank water on the day of the full marathon. In this study, three subjects were removed after exceeding the criteria for maximum levels of general biomarkers, including C-reactive protein and urinary albumin, in sedentary conditions before the full marathon.

**Table 1.** Participant characteristics

|  | Age | Height | Weight | Blood pressure<br>Diastolic/Systolic | 12 min<br>running | Marathon time |
| --- | --- | --- | --- | --- | --- | --- |
|  | (year) | (cm) | (kg) | (mmHg) | (m) | (h:m:s) |
| AVE | 25. 2 | 172. 0 | 64. 9 | 125. 7 / 74. 7 | 3181. 3 | 3:27:14 |
| SD | 7. 3 | 5. 3 | 6. 9 | 9. 1 / 7. 9 | 332. 2 | 0:58:48 |

### Experimental design

Measurements were collected immediately before (Pre) and after (Post) the full marathon, as well as two hours after completing the full marathon (2H), and the day after the full marathon (D1). The subjects drank only water between the Post and 2H measurements. The full marathon was completed at a temperature of 12.7 °C with a humidity level of 60.6%.

Blood and urine samples were collected early in the morning or after the full marathon. Blood samples were divided into plasma with heparin natrium and serum. Plasma samples were separated by centrifugation for 15 min at 4°C at 3000 rpm. Serum samples were separated by centrifugation for 15 min at 3000 rpm after sitting for 30 min at room temperature. These samples were stored at - −80°C until further analysis.

### Extracted cell-free DNA in blood and urine

Serum cfDNA (500 μL) was extracted using Serum Cell-Free Circulating DNA Purification Mini Kits (Norgen Biotek Corp.). Urinary cfDNA (10 mL) was extracted using Urine Cell-Free Circulating DNA Purification Mini Kits (Norgen Biotek Corp.). Extracted serum and urinary cfDNA were diluted in 30 μL and 50 μL of Elution Buffer, respectively.

Total cfDNA was measured using an Agilent Bioanalyzer 2100 (Agilent Technologies Corp.) using High-Sensitivity DNA kits, following the manufacturer’s instructions. The small fragment size in serum cfDNA was below 200 bp, while the large fragment size in cfDNA was above 1000 bp, as determined by previous studies^28^. The levels of each fragment size in the serum cfDNA were analysed in terms of peak value levels of small and large fragment sizes in the serum cfDNA. Other fragment sizes (201 bp-999 bp) in serum cfDNA were not analysed, because their conspicuous peak values were not observed.

### Biomarkers

Serum creatine kinase (CK)activity and myoglobin (Mb) levels were measured as skeletal muscle damage markers. Serum creatinine kinase, urinary amylase, albumin, and creatinine levels were also measured. Estimated glomerular filtration values (eGFRs) using serum creatinine concentrations were calculated by the Modification of Diet in Renal Disease (MDRD) equation for Japanese individuals. “eGFR (ml/min/1.73 m2) = 194 × [Concentration of serum CRE (mg/dl)]^−1.094^×[Age]^−0^’^287^”

White blood cells (WBCs) and their isoforms were measured as inflammation markers. Serum lactoferrin levels were measured using ELISA according to the manufacturer’s instructions. The analysis was conducted at three time points: Pre, Post, and 2H.

### Statistical analysis

All results are presented as mean or mean ± standard deviation. GraphPad Prism 7 software (GraphPad, Inc., La Jolla, CA, USA) was used. All data were analysed with non-parametric tests of homoscedasticity. All results were analysed using Dunn’s multiple comparisons method. Correlation analysis was conducted using the Spearman method. Significance levels were set at *p* < 0.05 or *p* < 0.01.

## Results

### Serum and urinary cfDNA concentrations

Serum cfDNA levels after a full marathon are shown in Fig. 1. Total serum cfDNA levels were increased immediately after the full marathon and showed higher values at 2H than at Pre, although serum cfDNA levels at 2H were lower than those at Post. Serum cfDNA small fragment levels were increased at Post compared to those at Pre. The levels of serum cfDNA large fragments were increased immediately at Post and remained high at 2H compared to those at Pre. Representative electropherogram results of serum cfDNA showed peak values from 170 to 180 bp after the full marathon. Values over 1000 bp were also present at 2H.

**Figure 1.**
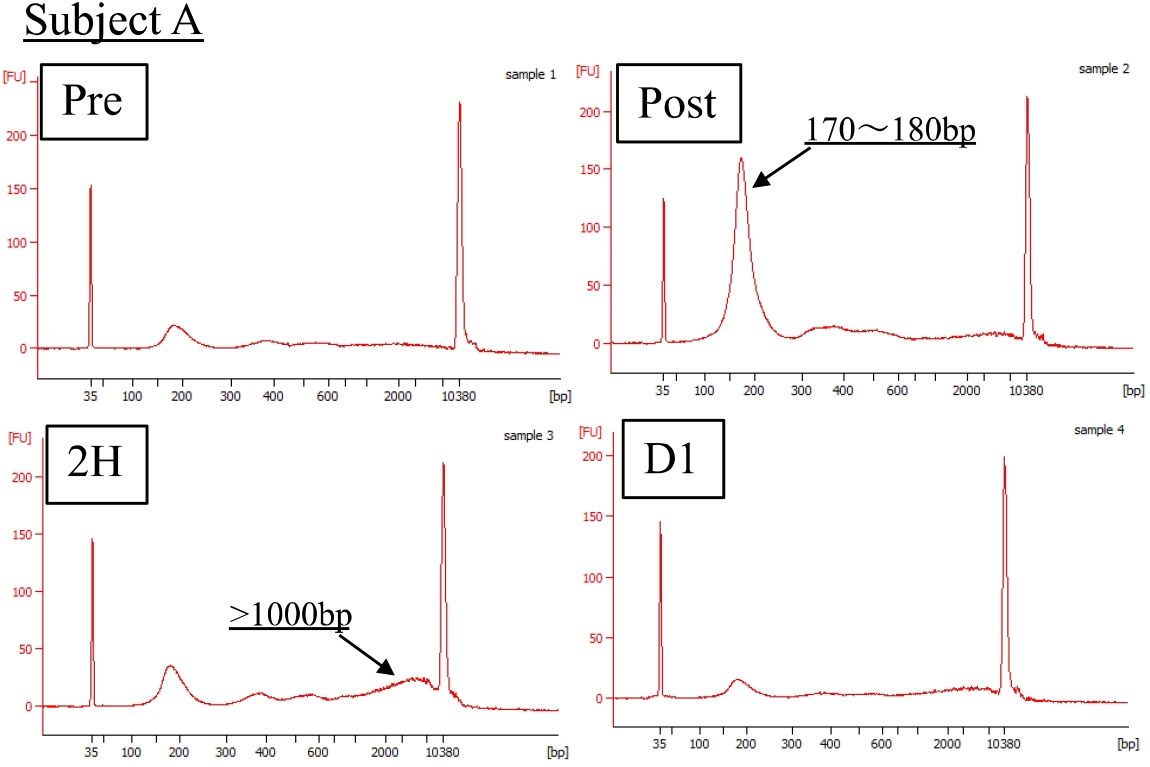
Serum cell–free DNA fragment size profiles in a full marathon Serum cell-free DNA fragment size profiles. Capillary electropherogram showing the fragment sizes of cfDNA extracted from serum samples after a full marathon by a representative subject.

Urinary cfDNA levels after the full marathon are shown in Fig. 2. Total urinary cfDNA levels were increased immediately after the full marathon. It was not possible to analyze the data pertaining to fragment size in urinary cfDNA due to the level placed on the marker. Representative electropherogram results for the urinary cfDNA showed peak values from 35 to 50 bp after the full marathon. Values over 1000 bp also appeared at Post and 2H.

**Figure 2.**
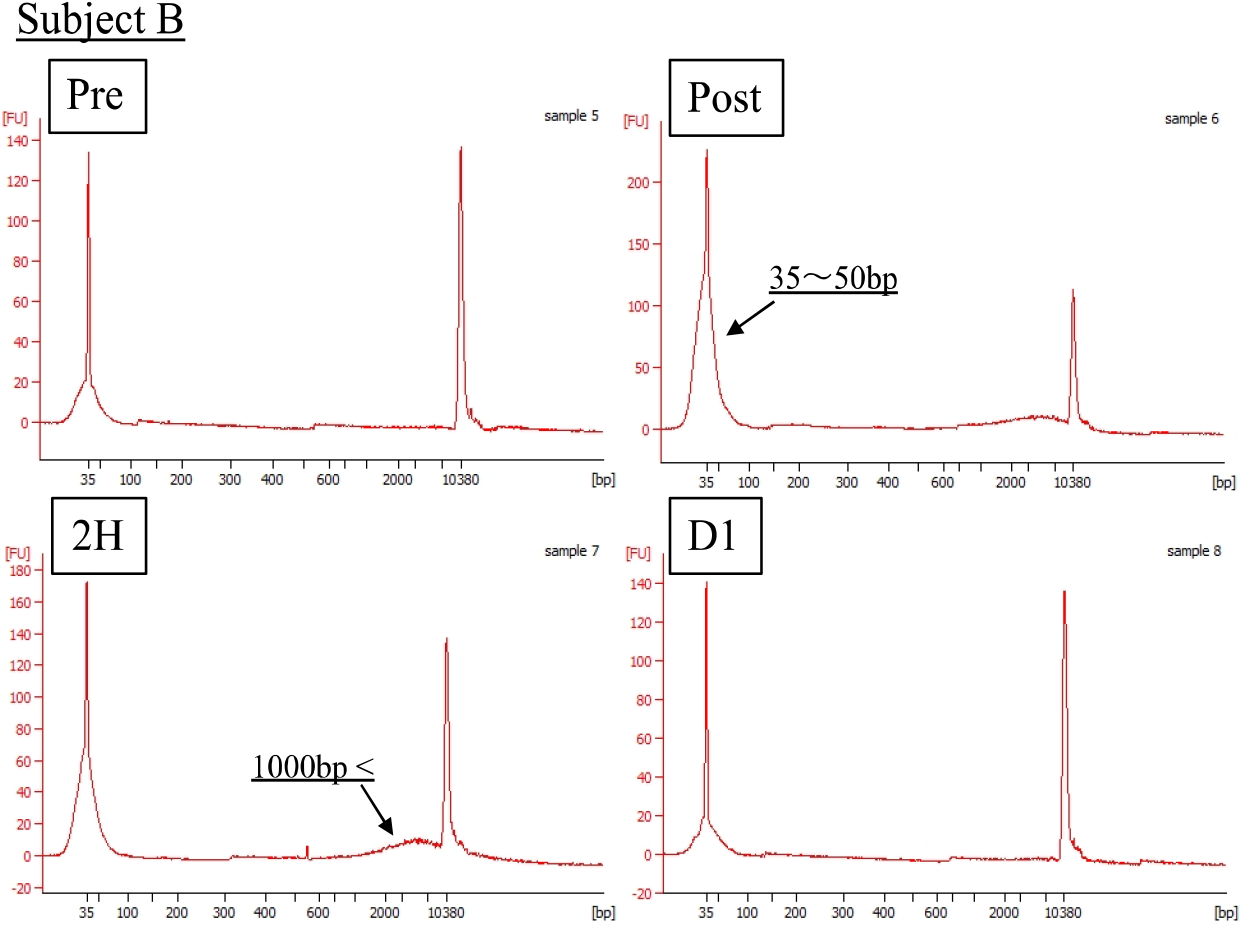
Urinary cell–free DNA fragment size profiles in a full marathon Urinary cell-free DNA fragment size profiles. Capillary electropherogram showing the fragment sizes of cfDNA extracted from a urinary sample after a full marathon by a representative subject.

### Biomarker profiles in blood and urine

Serum and urine biomarkers are shown in Table 2. Serum CK activity and Mb levels, the muscle damage markers, were increased after the full marathon, and maintained significantly higher values on D1 compared to those at Pre (*p* < 0.05, *p* < 0.01). Serum CK activity peaked on D1, while serum Mb levels peaked at Post or 2H. Serum creatinine levels were increased at Post and 2H compared to those at Pre. eGFR showed the same pattern of results as serum creatinine. Amylase levels in one of the urinary markers were unchanged after the full marathon. Urinary creatinine and albumin levels were increased after the full marathon (*p* < 0.01). Urinary albumin levels maintained a high value at 2H compared to those at Pre (*p* < 0.01). The albumin/creatinine ratio (ACR) showed the same results as that of urinary albumin (*p* < 0.01).

**Table 2.** Biomarkers after a full marathon

|  | Pre | Post | 2H | Day1 |
| --- | --- | --- | --- | --- |
| CK (U/L) | 151.6±44.3 | 824.5±455.5 * | 1394.7±801.5 ** | 3247.8±2570.7 ** |
| Mb | 30.6±7.9 | 2083.5±1372.9** | 1750.9±952.9 ** | 232.8±193.7 * |
| Creatinine | 0.86±0.11 | 1.04±0.19 ** | 0.96±0.15 * | 0.88±0.12 |
| eGFR (ml/min/1.73m²) | 92.4±13.0 | 76.5±15.2 ** | 83.3±14.7 * | 89.0±12.2 |
| Urinary amylase (U/L) | 381.1±157.4 | 494.0±321.6 | 426.5±289.0 | 424.5±178.0 |
| Urinary albumin (mg) | 3.3±1.3 | 36.6±38.0 ** | 16.6±16.1 ** | 4.6±2.3 |
| Urinary creatinine (mg/dL) | 200.1±65.2 | 283.2±161.8 ** | 223.0±156.1 | 224.5±84.2 |
| Urinary ACR | 16.8±6.3 | 122.3±111.5 ** | 84.8±74.4 ** | 21.3±8.4 |

CK, creatine kinase; Mb, myoglobin; ACR, albumin/creatinine ratio. \* $p < 0.05$ , \*\* $p < 0.01$ vs Pre

### WBC, platelet, and lactoferrin levels

The WBC, platelet, and lactoferrin results are shown in Fig. 3. WBCs and neutrophil levels peaked significantly at Post, and showed significantly high values at 2H and D1, despite the decrease from 2H (*p* < 0.05, *p* < 0.01). PLT was increased after the full marathon and maintained significantly higher values until 2H compared to those at Pre (*p* < 0.05, *p* < 0.01). Serum lactoferrin levels were significantly increased at Post and 2H compared to those at Pre (*p* < 0.01).

**Figure 3.**
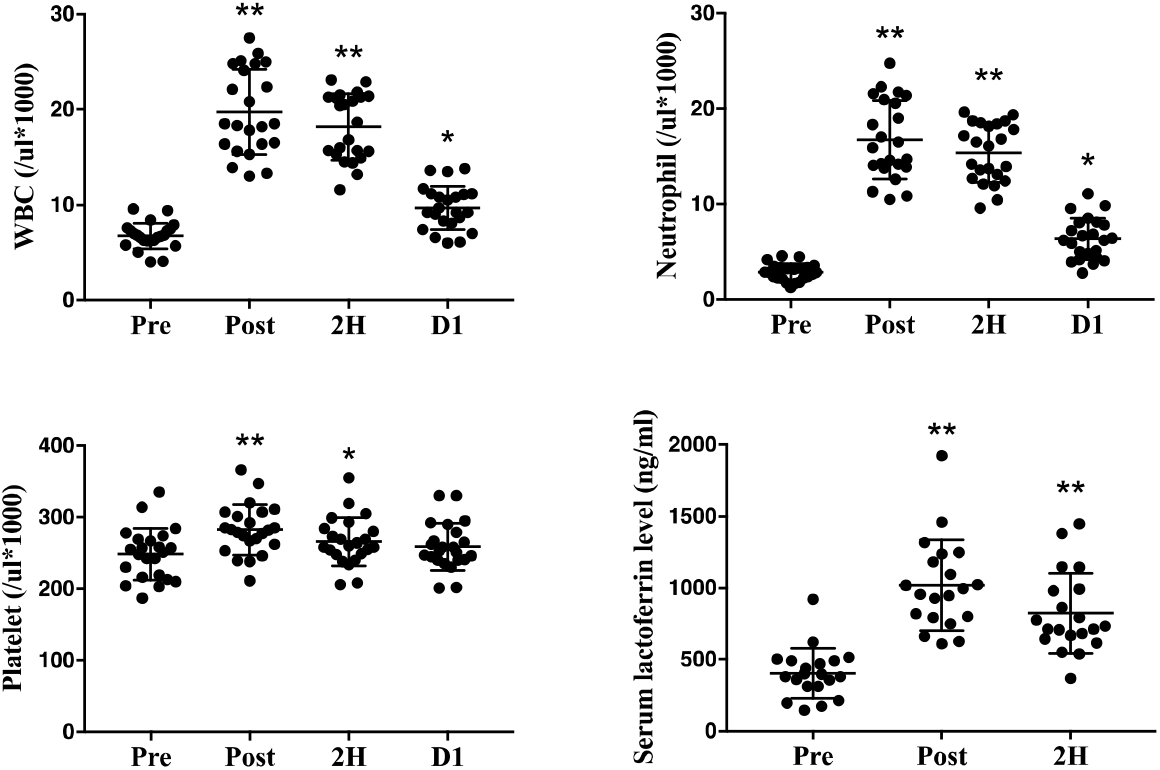
White blood cell counts and platelet and serum lactoferrin levels White blood cell counts and platelet and serum lactoferrin levels after a full marathon. Data are presented as mean ± SD and individual plots. p values indicate significant differences compared to Pre (\**p* < 0.05; ** *p* < 0.01). White blood cell counts and platelets, n = 23; serum lactoferrin levels, n = 20.

## Discussion

In this study we investigated the possibility of using a novel biomarker as an index for objectively describing body conditions and fatigue during exercise. Strenuous physical exercise, such as a full marathon, causes local damage to the muscle tissues. Muscle damage markers, such as serum CK activity and myoglobin levels, increase due to muscle damage^4,6–8,27^. In this study, the increased serum CK activity was delayed after the full marathon. These kinetics are in accordance with those of previous studies^7,16^. In contrast, serum myoglobin levels increased immediately at Post, and decreased one day later, but were significantly higher than at Pre. Investigation of the dynamics of these markers confirmed that the muscle tissues suffered damage due to the full marathon.

### Increased serum cfDNA and fragment size after a full marathon

Blood cfDNA levels transiently increase during various exercises, such as resistance and endurance exercises^11^. In addition to these exercise styles, this study suggests that serum cfDNA levels increase rapidly after a full marathon. This increase in cfDNA levels was immediately apparent, and then decreased to the baseline one day later. Our findings were in accordance with those of previous studies^12,15,16^. Levels of neutrophils and lactoferrins, which are the origin and component parts of NETs, increased after a full marathon^6,8^. Neutrophils induced during endurance exercise may function as host defences, involved in processes such as phagocytosis, degranulation, cytokine production, and, most recently described, NET production^29,30^. NET formation is considered to be the origin of increased cfDNA levels upon exercise^31^. On the other hand, copy number of cell free-mitochondria DNA (cf-mtDNA) also was increased by acute physical stress in previous study^32,33^. However, these results indicated copy number but not concentration in circulating blood. In addition, Hummel et al. reported copy number of cfDNA showed higher value than that of cf-mtDNA after acute exercise^32^. Therefore, these results suggested that almost all cfDNAs induced by a full marathon might have been derived from NETs.

In this study, the fragment size of serum cfDNA accumulated after a full marathon was measured. Our findings suggest that the cfDNA fragment size Pre marathon peaked at approximately 170 bp-180 bp (short fragments), and these short fragments accumulated immediately after the full marathon. cfDNA fragments >1000bp in size (large fragments) accumulated later, and the peak was lower than that of the short fragments. It has been reported that normal human plasma samples primarily contain cfDNA fragments of around 180 bp, and cfDNA fragments larger than 2000 bp cannot be detected. The primary origins of the circulating DNA in the blood are apoptotic, but not necrotic, cells^34^. In addition, small portions of the nucleus are released as NETs, a phenomenon which was most recently described in vivo after infection^35^. Therefore, cfDNA fragments shorter than 200 bp are probably derived from apoptosis, a form of programmed cell death, and possibly by NETosis, which releases NET formation from activated neutrophils. It has been found that plasma in cancer patients shows a time-dependent increase in cfDNA fragments over 10,000 bp in size. These larger fragments of cfDNA have been hypothesised to originate from the death of cells via necrosis^12^. A full marathon also induces necrosis through damage to the skeletal muscle^3^. Some of the subjects in this study also accumulated larger cfDNA fragments of over 1,000 bp. Short fragments were present immediately after a full marathon, whereas large fragments were induced 2 h later in most subjects. These results suggest that large fragments may be released by a different mechanism from short fragments, and contain DNA from the necrotic cells of damaged muscle tissues. In this way, differences in the timing of distribution and release of cfDNA fragments of different sizes may be due to their different origins and mechanisms. Confirmation of the distribution in cfDNA fragment size may make it possible to identify cfDNA derived from either type of cell death.

### Urinary cfDNA after a full marathon

This study showed for the first time, to the best of our knowledge, that urinary cfDNA levels were increased immediately after a full marathon. In the same way as serum samples taken after a full marathon, the levels of urinary cfDNA increased immediately after the exertion, and recovered to baseline levels one day later. Urinary cfDNA is derived either from cells shed into the urine from the genitourinary tract, or from cfDNA in systemic circulation passing through glomerular filtration^36,37^. The glomerular barrier discriminates between transported solutes based on size, charge, and shape^38^. In this study, ACR and eGFR, an index of renal function, showed significant increases after the full marathon. There has been a report of acute kidney injury developing after a full marathon^6^. Therefore, high-molecular-weight albumin was excreted through the kidney barrier into the urine, thereby increasing the levels of urinary cfDNA after the full marathon. Since marathon runners have been observed to develop a transient acute kidney injury (AKI) with urine sediment in a previous study, the increase in serum creatinine and urinary ACR after a full marathon may also represent a kidney injury^6,39^. It is possible that renal condition and urinary cfDNA levels are strongly related, as it has also been reported that AKI increases urinary cfDNA levels ^40^.

Urinary cfDNA fragment size, as well as serum cfDNA fragment size, changed after a full marathon, although the sizes were different. The length of serum cfDNA fragments increased by approximately 180 bp, whereas urinary cfDNA fragments increased by approximately 35 bp-50 bp in size. The primary function of the glomeruli in the kidneys is to filter low molecular weight wastes into the urine, and regulate the passage of albumin and large macromolecules, which are necessary for homoeostasis^38^. Large fragments in urinary cfDNA accumulated late, as was seen in serum cfDNA. If the large fragments in serum cfDNA are derived from necrotic cells, it is likely that they are also derived from the same source in urine. The large fragments derived from necrotic cells may arise from the urinary tract through the kidney filter, or from muscle damage due to exercise. It is possible that the large fragments in urinary cfDNA are related to the muscle damage markers, but this study did not measure urinary muscle damage markers. In a future study, it will be necessary to measure urinary muscle damage markers as well as urinary cfDNA level.

In conclusion, serum and urinary cfDNA levels were transiently increased after a full marathon. As regards size profile of cfDNA, the short fragment size of serum cfDNA accumulated immediately after the full marathon and large fragments accumulated later. Urinary cfDNA also had fragments of larger size after a full marathon. However, it is necessary to investigate the effects of intensity, time, style of exercise, and sex on cfDNA levels to use this molecule an exercise biomarker, as its physiological significance remains to be clarified.

### Perspective

Sports medicine research into exercise biomarkers aims to improve health, performance, and recovery. However, there are few recommendations for biomarkers for tracking changes in individuals participating in physical activity and exercise training programs^41^. Previous studies have reported that aerobic and anaerobic exercise induces an immediate and transient increase in blood cfDNA levels^11^. The present study suggested that blood cfDNA levels peaked immediately after a full marathon. Urinary cfDNA levels also increased transiently. It is clinically important to identify immediate, non-invasive biomarkers such as cfDNA for the development of tailored exercise programs to prevent overwork and disease. Haller et al. (2018) reported that cfDNA showed the most pronounced increase that has ever been reported, compared to other biomarkers, in an acute intermittent exercise setting^13^. The results of this study provide new possibilities for the early detection and monitoring of internal stress caused by exercise.

## Authorships

K Tokinoya, YA, and K Takekoshi contributed to the design. YS, K Tokinoya, NS, TS, YY, KK, SK, KI, TK, YN, SM, and K Takekoshi performed data acquisition. YS, K Tokinoya, TS, and K Takekoshi analysed and interpreted the data. YS and K Tokinoya performed the statistical analyses. YS and K Tokinoya drafted the manuscript. All authors supervised and edited the manuscript. All authors have reviewed the manuscript. All authors provided final approval of this version of the manuscript for publication and agreed to be accountable for all aspects of the work.

## Acknowledgments

All subjects participated in this study. In addition, we would like to thank Editage (www.editage.com) for English language editing. This work was supported by JSPS KAKENHI (Grant Number: 19K11434). There are no conflicts of interest to disclose. The results of the study are presented clearly, honestly, and without fabrication, falsification, or inappropriate data manipulation.

